# Sex differences in migraine: A twin study

**DOI:** 10.1101/2021.10.27.466155

**Authors:** Fitzgerald Morgan, Ursula G. Saelzler, Matthew S. Panizzon

## Abstract

Migraine is a neurological disorder with a prominent sex difference such that two thirds of sufferers are female. The mechanisms behind the preponderance of migraine in women have yet to be elucidated. With data on 51,872 participants from the Swedish Twin Registry, we report results from two distinct analyses intended to clarify the degree to which genetic and environmental factors contribute to sex differences in migraine. First, we fit a sex-limitation model to determine if quantitative genetic differences (i.e., is migraine equally heritable across men and women) and/or qualitative genetic differences (i.e., are different genes involved in migraine across men and women) were present. Next, we used a multilevel logistic regression model to compare the prevalence of migraine in individuals from opposite-sex and same-sex twin pairs to determine whether differences in the prenatal hormone environment contribute to migraine risk. In the final analytic sample, women were found to have a significantly higher rate of migraine without aura relative to men (17.6% vs 5.5%). The results from an ADE sex-limitation model indicate that migraine is equally heritable in men and women, with a broad sense heritability of .45, (95% CI = .40 - .50), while results from a reduced AE sex-limitation model provide subtle evidence for differences in the genes underlying migraine across men and women. The logistic regression analysis revealed a significant increase in migraine risk for females with a male co-twin relative to females with a female co-twin (OR = 1.51, 95% CI = 1.26 – 1.81). These results suggest that the prominent sex difference in migraine prevalence is not entirely accounted for by genetic factors, while demonstrating that masculinization of the prenatal environment may increase migraine risk for females. This effect points to a potential prenatal neuroendocrine factor in the development of migraine.

## 1 INTRODUCTION

Migraine is a severe neurological disease characterized by repeated transient symptomatic episodes, affecting approximately 1 in 7 people. Although migraine presents with varying degrees of severity, it is the second largest cause of disability worldwide, and in 2020 the economic burden in the U.S. was estimated at $78 billion (GBD, 2019; Vos et al., 2017; Polson et al., 2020). A prominent sex difference in migraine exists such that roughly 18% of adult women report experiencing migraine compared to only 6% of adult men (Burch, Rizzoli & Loder, 2018; Victor et al., 2010; Sacco, Ricci, Degan & Carolei, 2012; Lipton et al., 2001). While the preponderance of migraine in women is striking, to date little is known regarding the origins of this sex difference. Analytic methods available in the context of the classical twin design offer unique opportunities to explore this issue.

The classical twin design relies on the comparison of monozygotic (MZ) twins, who are genetically identical, with dizygotic (DZ) twins, who share on average 50% of their segregating DNA, to make inferences regarding the degree to which genetic and environmental factors contribute to disease risk within a population. As a variation of the classical twin model, the sex-limitation model employs data from male and female same-sex (SS) MZ and DZ twins, as well as opposite-sex (OS) DZ twins to test for qualitative and quantitative genetic differences between men and women (Neale & Cardon, 2013). *Qualitative* genetic differences assess whether different genes are involved in disease risk across the sexes, while *quantitative* genetic differences determine if disease risk is equally heritable across the sexes.

Twin and family studies have shown that migraine has a substantial genetic component. First degree relatives of individuals with migraine have a 1.4 to 3.0 times higher risk of experiencing the disease compared to those with no immediate family history of migraine (Russell, Iselius & Olesen, 1995; Low, Cui & Merikangas, 2007). Concordance rates for migraine are also consistently higher among MZ twins relative to DZ twins, and twin-based heritability estimates range from .34 to .57, indicating that 34 to 57% of the risk liability (i.e., variation in migraine risk within a population) is attributable to genetic factors (Ulrich, Gervil, Kyvik, Olesen & Russell, 1999; Cox, et al., 2012; Wang, Lui & Zhao, 2008; Nyholt et al., 2004; Ziegler et al., 1998).

Several twin studies have explored the possibility of sex differences in the genetic and environmental determinants of migraine. In a study of over 8,000 Finish twin pairs, the heritability of migraine was essentially equal between men and women (.42 and .47, respectively), suggesting little quantitative difference in the magnitude of genetic effects (Honkasalo, Kapiro, Winter, Heikkila, Sillanpaa & Koskenvuo, 1995). These findings were largely replicated in a subsequent study of 5,000 Danish twins (Gervil, Kaprio, Olsen & Russell, 1999). In one of the largest twin studies conducted, Mulder et al. (2003) examined the prevalence and heritability of migraine in eight twin cohorts across six countries (n = 330—12,121) and assessed both quantitative and qualitative differences. Heritability estimates ranged from .33 to .56 and could be equated between men and women both within and between countries, suggesting no quantitative sex differences. Similarly, looking at the *individual*-cohort level, there was no evidence for qualitative sex differences

Although Mulder et al. provided strong evidence for a lack of quantitative genetic difference between men and women, the analysis did not leverage the larger combined sample in the test of qualitative genetic differences. One potential reason for this could be because the definition of migraine ranged from a single item response regarding “headache attacks” or receiving a physician diagnosis, to detailed surveys that aligned with the International Headache Society’s (IHS) diagnostic criteria. Thus, a large-scale, high-powered test of qualitative genetic differences in a population with a detailed classification of migraine has yet to be conducted. The first goal of the present study was to replicate the sex-limitation model investigation done by Mulder et al. in a larger sample with greater statistical power, and for which migraine has been diagnosed using a consistent, detailed set of criteria.

In addition to the sex-limitation model, the comparison of prevalence rates of disease in twins from OS DZ pairs relative to SS pairs through an opposite-sex twin comparison paradigm provides insight into how variation within the prenatal environment can impact brain development and disease risk. While not a genetically informative analysis, this design does allow for inferences to be made regarding the role of the prenatal hormone milieu on lifetime outcomes.

There is substantial evidence to suggest that sex steroid hormones may contribute to the observed sex difference in the prevalence of migraine. The ‘estrogen withdrawal hypothesis’ (Reddy et al., 2021) specifically suggests that changes in serum estradiol levels are capable of precipitating migraine attacks. In support of this hypothesis, before puberty, when sex steroid hormones are largely dormant, the prevalence of migraine is equal in boys and girls (Lipton & Bigal, 2005; MacGregor, 1997). In women, age at menarche is associated with age at migraine onset such that each one-year delay in onset of menarche decreases the odds of migraine by seven percent (Maleki, Kurth & Field, 2017). During the reproductive period, menses, which corresponds to a cyclical decline in estradiol and progesterone, is associated with an elevated risk of migraine attack (Johannes et al., 1995; Stewart et al., 2000; Wöber et al., 2007). During perimenopause, a transition period characterized by the fluctuation and progressive loss of estradiol, migraine attacks in women commonly increase in frequency (Martin et al., 2016; Wang et al., 2003). Finally, a lower incidence of migraine is observed following the menopausal transition (Sacco, Ricci, Degan & Carolei., 2012; Martin et al., 2016; Wang et al., 2003). Migraine throughout the male lifetime is understudied; however, men with migraine have been found to have lower testosterone and higher estradiol levels relative to age matched controls (Shields, Seifert, Shelton, & Plato, 2019; van Oosterhout et al., 2019).

In conjunction with the potential influence of circulating sex steroid hormones, evidence from both animal and human twin studies suggest an association between the prenatal hormonal milieu and disease risk later in life. Expressly, the ‘twin testosterone transfer hypothesis’ posits that females gestated with a male co-twin will demonstrate physiological and behavioral masculinization (Tapp, Mayberry & Whitehouse, 2011). In support of this hypothesis, several rodent studies have shown that, compared to female rodents gestated with other females, female fetuses gestated with males demonstrated higher blood concentrations of testosterone and lower levels of estradiol, increased anogenital distance, and more aggressive and territorial behaviors (Ryan & Vandenbergh, 2002; Breedlove, 1994; Tapp, Maybery, & Whitehouse, 2011). Critically, prenatal administration of an anti-androgen prevented the masculinization of female rodents, suggesting that the effects were driven by increased androgen exposure in utero. In humans, twin studies have demonstrated that females with a male co-twin exhibit lasting masculinization of perception, cognition, personality and risk of dementia (Tapp, Maybery, & Whitehouse, 2011; Loehlin & Martin, 2000; Vuoksimaa et al., 2010; Verweij, Mosing, Ullen & Madison, 2016; Luo et al., 2020). Therefore, the second goal of this study was to conduct a novel comparison of SS and OS twins to examine the role of the prenatal environment, specifically hormonal variation within the prenatal environment, on migraine risk.

In the present study, we utilized data from the Swedish Twin Registry (STR), one of the largest population-based twin registries in the world (Lichtenstein et al., 2006), to conduct a comprehensive, genetically informed examination of sex differences in migraine. With data from over 18,000 complete twin pairs, this study represents the largest investigation of qualitative genetic differences in migraine ever conducted. In addition, we leveraged the presence of OS twin pairs to test the novel hypothesis that the prenatal hormonal milieu contributes to migraine risk in women. We hypothesized that the presence of a male co-twin would decrease migraine risk in females due to androgenization effects produced by the inter-twin transfer of testosterone.

## METHODS

### Participants

The sample was pooled from 70,223 SS and OS MZ and DZ twins who participated in the STR. Structured interviews evaluating multiple health outcomes, including migraine, were administered as part of two independent studies: the Screening Across the Lifespan Twin Study (SALT) questionnaire one and two, and the Study of Twin Adults – Genes and Environment (STAGE). The analytic sample contains data from 51,872 twins with complete data born between 1959 and 1985 including 18,365 SS female twins, 15,335 SS male twins, 8,433 OS male twins and 9,739 OS female twins. See demographic information for the analytic sample in Table 1.

**Table 1.**
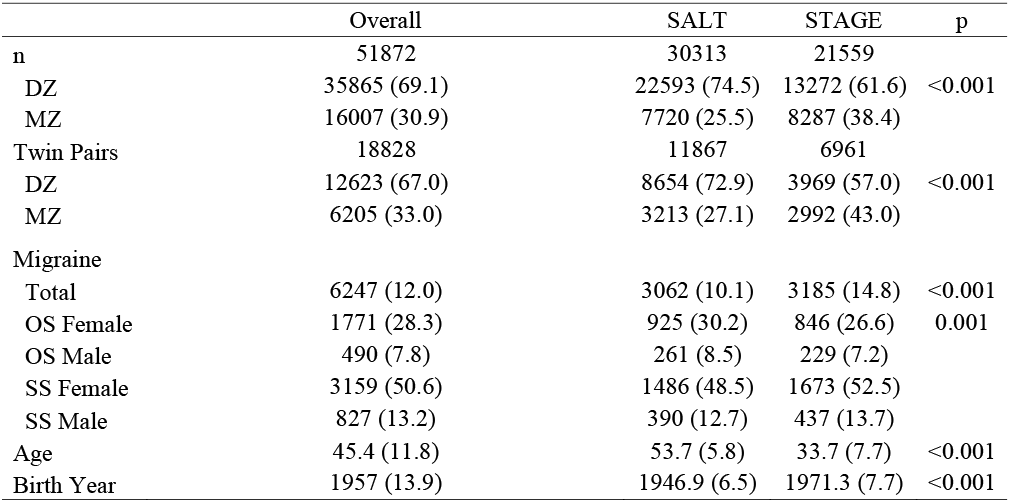
Demographics by twin pair type. DZ: Dizygotic, MZ: Monozygotic, OS: Opposite-sex; SS: Same-sex. SALT: Screening Across the Lifespan Twin Study, STAGE: Study of Twin Adults – Genes and Environment, Age reflects age at survey. Counts shown as: number (column %), continuous variables shown as mean (SD). P-values reflect comparison of SALT and STAGE cohorts.

|  | Overall | SALT | STAGE | p |
| --- | --- | --- | --- | --- |
| n | 51872 | 30313 | 21559 |  |
| DZ | 35865 (69.1) | 22593 (74.5) | 13272 (61.6) | <0.001 |
| MZ | 16007 (30.9) | 7720 (25.5) | 8287 (38.4) |  |
| Twin Pairs | 18828 | 11867 | 6961 |  |
| DZ | 12623 (67.0) | 8654 (72.9) | 3969 (57.0) | <0.001 |
| MZ | 6205 (33.0) | 3213 (27.1) | 2992 (43.0) |  |
| Migraine |  |  |  |  |
| Total | 6247 (12.0) | 3062 (10.1) | 3185 (14.8) | <0.001 |
| OS Female | 1771 (28.3) | 925 (30.2) | 846 (26.6) | 0.001 |
| OS Male | 490 (7.8) | 261 (8.5) | 229 (7.2) |  |
| SS Female | 3159 (50.6) | 1486 (48.5) | 1673 (52.5) |  |
| SS Male | 827 (13.2) | 390 (12.7) | 437 (13.7) |  |
| Age | 45.4 (11.8) | 53.7 (5.8) | 33.7 (7.7) | <0.001 |
| Birth Year | 1957 (13.9) | 1946.9 (6.5) | 1971.3 (7.7) | <0.001 |

### Assessment of migraine without aura

Sufficient data from both SALT and STAGE allowed for the utilization of the IHS classification of migraine without aura (MO). Items selected best adhered to the IHS International Classification of Headache Disorders version three (ICHD-3) MO criteria A, B, C and D including attack recurrence, duration, characteristics (unilateral location, pulsating quality, moderate or severe pain intensity, aggravation by or causing avoidance of routine physical activity), and symptoms (nausea and/or vomiting, photophobia and phonophobia; Table 2). Notably, STAGE did not have a proxy for duration of attack (between four and 72 hours), so this criterion was omitted from migraine classification for the STAGE cohort.

**Table 2.** Migraine diagnostic criteria and associated questionnaire items. The International Classification of Headache Disorders Third Edition (ICHD-3) migraine without aura criteria for diagnosis as compared to the variables used from the Screening Across the Lifespan Twin Study (SALT) and the Study of Twin Adults – Genes and Environment (STAGE) databases.

| ICHD-3 Migraine Without Aura Criteria |  | SALT | STAGE |
| --- | --- | --- | --- |
|  |  | Question | Question |
| A | **At least 5 attacks fulfilling criteria B-D** | Have you or have you ever had recurrent headaches, which did not come from a cold, fever or hangover? | Have you ever had recurrent headaches i.e., headaches that recur more or less often with headache-free intervals in between? |
| B | **Attacks lasting 4-72h (untreated or treated)** | How long does your headache usually last without medication or if the medication did not help? | NA |
| C | *At least two of the following characteristics:* |  |  |
|  | **Unilateral location** | Is the headache mostly on * one side * of the head, and then behind the eye, in the temple or forehead? | Is the headache located to one side, either left or on right but not both sides at the same time? |
|  | **Pulsating quality** | Pulsating / throbbing-type-of-ache? | Does the headache feel pounding or pulsating? |
|  | **Moderate or severe pain intensity** | How would you describe your headache? [Intensity rating] | Is the headache severe? |
|  | **Aggravation by or causing avoidance of routine physical activity** | Is the headache aggravated by normal physical activity, for example when you walk up stairs? | Does the headache get worse by moderate physical activity (e.g., walking, climbing stairs) or by coughing and sneezing? |
| D | *During attack – at least one of the following:* |  |  |
|  | **Nausea and/or vomiting** | Do you usually feel sick during the headache? | Is the headache accompanied with nausea? |
|  |  | Do you usually vomit during the headache? | Is the headache accompanied vomiting? |
|  | **Photophobia and phonophobia** | During the headache, do you often get an increased sensitivity to light? | Is the headache accompanied with increased sensitivity to light? |
|  |  | During the headache you often get an increased sensitivity to sound? | Is the headache accompanied with increased sensitivity to sound? |
| E | **Not better accounted for by another ICHD-3 diagnosis** | NA | NA |

### Statistical analyses

#### Sex-Limitation Model

Within the classical twin design, data from MZ and DZ twins are used to decompose the variance of a trait/phenotype into **additive genetic (A) influences** (i.e., genetic effects attributable to multiple genes that exert their independent influence in a linear fashion), **non-additive genetic (D) influences** (i.e., genetic effects attributable to dominance and/or epistasis; Rettew et al., 2008), **common or shared environmental (C) influences** (i.e., environmental factors that make members of a twin pair similar to one another), and **unique environmental (E) influences** (i.e., environmental factors that make members of a twin pair different from one another, including measurement error; Neale and Cardon, 2013).

Additive and non-additive genetic influences are assumed to correlate perfectly (r_A_ and r_D_ = 1.0) between MZ twins because they are genetically identical. In contrast, DZ twins share on average 50% of their segregating DNA and are therefore assumed to demonstrate a correlation of r_A_ = .50 for additive genetic influences and r_D_ = .25 for non-additive genetic influences. When standardized, the combined additive and non-additive genetic influences provide an estimate of broad-sense heritability. The shared environment, by definition, correlates perfectly between both members of a twin pair, regardless of their zygosity. Unique or non-shared environmental influences are uncorrelated among twin pairs.

Due to issues of model identification, A, D, C, and E variance components cannot be simultaneously fit with only twin data; therefore, researchers are restricted to fitting either an ACE or ADE model. For the present study, examination of cross-twin tetrachoric correlations revealed that for both male and female SS twins the MZ correlation was more than twice that of the DZ correlation (Male Twins: r_MZ_ = .46, r_DZ_ = .19; Female Twins: r_MZ_ = .44, r_DZ_ = .11), indicating a minimal impact of shared environmental influences (C) and that an ADE model would provide the most appropriate representation of the data. Therefore, we utilized the ADE model within the sex-limitation model framework.

The sex-limitation model involves the application of the ADE model to data from male and female twins simultaneously, thereby allowing tests of heterogeneity of the variance components between the sexes (i.e., are the magnitudes of genetic and environmental contributions equivalent; see Figure 1). Specifically, *quantitative* sex differences are tested by constraining the genetic and environmental variance components to be equal across the male and female groups (i.e., A_M_=A_F_, D_M_=D_F_, E_M_=E_F_). Should these constraints result in a significant change in model fit (p < .05) relative to an unconstrained model, quantitative differences can be inferred. Furthermore, with the inclusion of OS twins, the sex-limitation model allows for estimation of the genetic correlation between male and female twins in OS pairs, which enables the additional test of *qualitative* genetic differences. Qualitative genetic differences are inferred when the genetic correlation between OS DZ twins deviates significantly from the value assumed for SS DZ twins (.5 for additive genetic influences and .25 for non-additive genetic influences).

**Figure 1.**
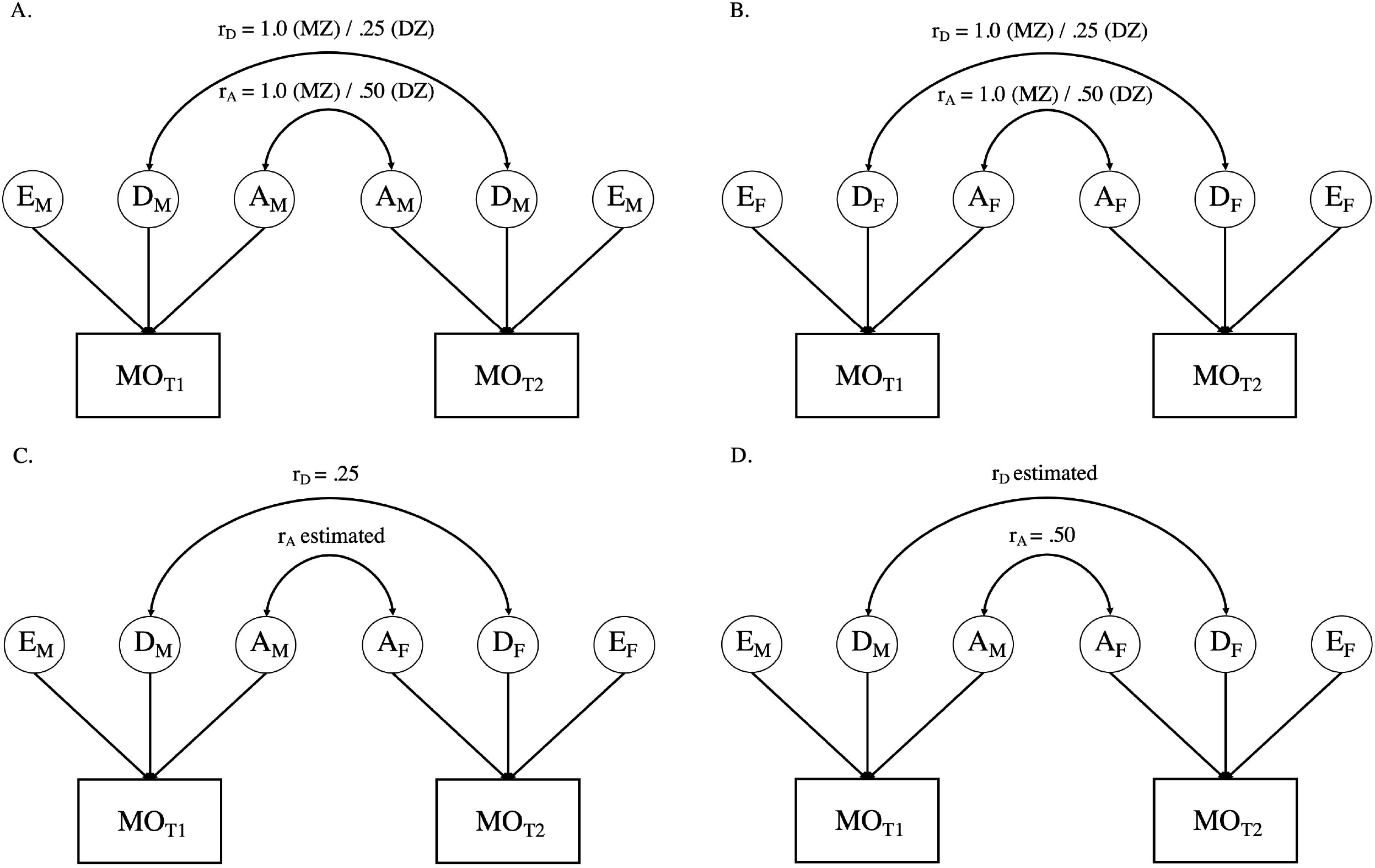
Panels A presents the path diagram for male-male twin pairs. Panel B presents the path diagram for female-female twin pairs. Panels C and D offer two variations of estimating variation in opposite sex twins. As noted in the text, due to issues of model identification, r_A_, and r_D_ cannot be simultaneously estimated. Variance components with the same subscript are constrained to be equal within and across all twin pairs. MO = Migraine without aura; T1 = twin 1; T2 = twin 2; A = Additive genetic influences; C = Common environmental influences; E = Unique or non-shared environmental influences. Aspects of the model with an “M” subscript indicate those elements that are specific to males, whereas those with an “F” are specific to females.

To evaluate all aspects of the sex-limitation model, a series of four models were fit and tested against the Saturated Model (see Table 3). The Saturated Model provides a mathematical representation of the data against which all theoretical (i.e., sex-limitation) models are compared. Model 1 addressed *qualitative* differences by allowing the correlation between the additive genetic components (r_A_) of the OS twins to be estimated while constraining the correlation of the non-additive genetic influences to the expected DZ twin value (i.e., r_D_ = .25; see Figure 1 Panel C). Model 2 inverted these constraints (restricting the additive genetic correlation to r_A_ = .5, in line with the expected DZ twin correlation and estimating the non-additive genetic correlation (rD); see Figure 1 Panel D). Model 3 constrained both the additive and non-additive genetic correlations to the expected DZ values (r_A_ = .5 and r_D_ = .25). While Models 1—3 allowed the ADE variance parameters to be independently estimated for men and women, Model 4 constrained the variance components to be equivalent across men and women (i.e., A_M_=A_F_, D_M_=D_F_, E_M_=E_F_) while maintaining the restraints on the additive and non-additive genetic correlations from Model 3, thereby allowing for the test of *quantitative* sex differences.

**Table 3.**
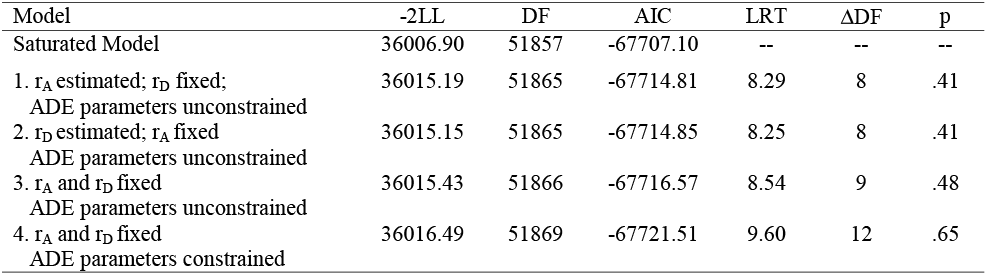
ADE sex-limitation model fitting results. All model fits are reported relative to the Saturated Model. −2LL = Negative 2 log-likelihood; DF = Degrees of Freedom; AIC = Akaike’s Information Criterion; LRT = Likelihood ratio test (differences in −2LL between models).

| Model | -2LL | DF | AIC | LRT | ΔDF | p |
| --- | --- | --- | --- | --- | --- | --- |
| Saturated Model | 36006.90 | 51857 | -67707.10 | -- | -- | -- |
| 1. $r_A$ estimated; $r_D$ fixed;<br>ADE parameters unconstrained | 36015.19 | 51865 | -67714.81 | 8.29 | 8 | .41 |
| 2. $r_D$ estimated; $r_A$ fixed<br>ADE parameters unconstrained | 36015.15 | 51865 | -67714.85 | 8.25 | 8 | .41 |
| 3. $r_A$ and $r_D$ fixed<br>ADE parameters unconstrained | 36015.43 | 51866 | -67716.57 | 8.54 | 9 | .48 |
| 4. $r_A$ and $r_D$ fixed<br>ADE parameters constrained | 36016.49 | 51869 | -67721.51 | 9.60 | 12 | .65 |

Models were fit to all available raw data by full information maximum-likelihood, using the structural equation modeling software OpenMx (v2.19.5) within R (Boker et al., 2011; Neale et al., 2016). Given the binary nature of the outcome (i.e., positive or negative history of MO), threshold liability models were utilized whereby a threshold for migraine was estimated for male and female twins, and the phenotypic variance was constrained to 1.0 (Neale and Cardon, 2013). Evaluation of model fit employed the likelihood-ratio chi-square test (LRT), which is calculated as the difference in the negative 2 log-likelihood (−2LL) of a model relative to that of a comparison model. The Akaike’s information criterion (AIC) was used as a secondary indicator of model fit, with lower AIC values indicating a preferred balance between model fit and parsimony (Akaike, 1987). Significance of individual model parameters (e.g., r_A_, r_D_, A, D, E) was determined by calculation of likelihood-based 95% confidence intervals (CIs).

#### Opposite-Sex Twin Comparison

To compare the rates of migraine in SS and OS twin pairs, a multilevel logistic regression model was fit to all available data using the *glmer* function from the lme4 package (version 1.1.27.1; Bates, Maechler, Bolker & Walker, 2015) in R (version 4.0.4; R Core Team, 2021). Fit was accomplished by maximum-likelihood estimation via the Laplace approximation with the nlminb optimizer wrapped through optimx (Nash & Varadhan, 2011; Nash, 2014). Migraine diagnosis was included as the binary outcome (yes/no), predicted by study (included as a 3-level categorical variable: SALT questionnaire 1, SALT questionnaire 2 and STAGE), and twin-pair type (included as a 4-level categorical variable: SS female, SS male, OS male, and OS female, with SS female as the reference level). The SALT study utilized two versions of its medical history interview, and while the items were equivalent, the different structure of the assessments warranted distinct coding. Twin pair was included as a random effect to account for non-independence of the observations. Given that the primary aim of this analysis was to compare the prevalence of migraine among SS versus OS individuals as a function of sex (i.e., SS females versus OS females, and SS males versus OS males), an analogous model was fit to the data using SS male as the reference level.

## 2 RESULTS

Of the 70,223 STR participants, 1,525 individuals did not have information available regarding co-twin sex, and an additional 16,826 individuals did not have sufficient information to determine migraine status, resulting in a total analytic sample of 51,872 individuals, representing 18,828 complete twin pairs. Out of the analytic sample, 6,247 (12.0%) individuals met the diagnostic criteria for migraine. Of those meeting the diagnostic criteria, 79.0% were female, such that the prevalence of migraine was 17.6% for women and 5.5% for men.

### Qualitative and quantitative genetic difference

As shown in Table 3, both variants of the unconstrained ADE sex-limitation model (Models 1 and 2), provided good fits relative to the Saturated Model (i.e., both models resulted in a non-significant change in −2LL). Indeed, both models provided near identical fits and resulted in AIC values that were negligibly different from one another (−67714.81 vs. −67714.85). Liability thresholds for men and women were allowed to vary, as equating them resulted in a significant reduction in fit (p < .001), consistent with the substantially higher prevalence of migraine in women than men in the sample. Fixing the OS DZ twin genetic correlations to their expected values (Model 3: r_A_ = .50, r_D_ = .25) also resulted in a good fit relative to the Saturated Model (LRT = 8.54, ΔDF = 9, p = .48). Further, comparison of Model 3 with the variants of the unconstrained ADE sex limitation models (Models 1 and 2) indicated that constraining the genetic correlations did not result in a significant change in fit (fit relative to Model 1: LRT = 0.24, ΔDF = 1, p = .62; fit relative to Model 2: LRT = 0.28, ΔDF = 1, p = .59). The additional constraints did not significantly reduce model fit, which suggests that there is no *qualitative* sex difference in the genetic determinants of migraine (i.e., we found no evidence that migraine risk is driven by different genes in men and women).

Constraining the genetic and environmental variance components (A, D, E) to be equal across men and women (Table 3, Model 4) resulted in a non-significant change in fit relative to the Saturated Model and resulted in the lowest AIC value of all models tested. Change in fit relative to Model 3 (which tested for qualitative sex differences) was also non-significant (LRT = 1.06, ΔDF = 3, p = .79). Thus, here there is no evidence supporting *quantitative* genetic difference in migraine across the sexes.

In this final model, the broad sense heritability (A + D) of migraine was found to be .45 (95% CI: .40 – .50), indicating that 45% of the variation in risk for migraine was due to genetic factors. Much of this effect was attributable to non-additive genetic factors (D = .38, 95% CI: .18 - .50), with a relatively small contribution of additive genetic factors (A = .07, 95% CI: .04 - .26). The remainder of risk for migraine was accounted for by unique environmental factors (E = .55, 95% CI: .50 - .60).

#### Reduced Genetic Model

The presence of both additive and non-additive genetic influences complicated the test of qualitative sex differences as we were unable to simultaneously estimate r_A_ and r_D_ in the OS DZ twins. We therefore fit a series of three reduced genetic models to determine if simplifying the number of model parameters improved our ability to detect qualitative genetic differences. For all reduced models, non-additive genetic influences were constrained to zero (D = 0), resulting in a model where only additive genetic (A) and unique environmental (E) variance components were estimated.

The unconstrained AE sex-limitation model resulted in a non-significant change in fit relative to the Saturated Model (LRT = 17.24, ΔDF = 10, p = .07). It is worth noting, however, that the AIC value for the unconstrained AE sex-limitation model was larger than the AIC values for either of the ADE variants, highlighting the importance of non-additive genetic factors to migraine. Constraining the additive genetic correlation (r_A_) to .50 within the OS DZ twins resulted in a significant change in fit relative to the Saturated Model (LRT = 21.61, ΔDF = 11, p = .03) and the unconstrained AE sex-limitation model (LRT = 4.37, ΔDF = 1, p = .04). Thus, the estimated r_A_ of .32 (95% CI: .16 - .49) was found to significantly deviate from the assumed .50 value, indicating *qualitative sex differences in the additive genetic components of migraine between men and women*. The final best-fitting model under this reduced genetic influence scenario was obtained by constraining the A and E variance components to be equal across men and women (A = .42, 95% CI: .37 – .47; E = .58, 95% CI: .53 – .63), while also estimating the genetic correlation between OS twins at .32. This simplified model suggests that there are slight *qualitative* genetic differences in migraine across the sexes.

### Opposite sex twin comparison results

Both study and twin-pair type were significant predictors of migraine prevalence, as evidenced by significant improvements in fit compared to a pair of models with each predictor dropped (study: LRT = 40.7 ΔDF = 2, p < .001 and twin-pair type: LRT = 1542.9, ΔDF = 3, p < .001). Specifically, STAGE was associated with an increased odds of migraine (OR = 1.74, 95% CI = 1.46 – 2.07) compared to SALT questionnaire 1, while the two SALT cohorts did not differ (SALT questionnaire 1 OR = 1.07, 95% CI = 0.80 – 1.44). This difference is likely driven by the inability of the STAGE interview to address ICHD-3 criteria B (Table 2), resulting in a higher prevalence of migraine (14.8% versus 10.5% and 10.0% in SALT questionnaire 1 and 2, respectively).

As anticipated, due to the significantly higher prevalence of migraine among women than men, in comparison to SS females, both SS males and OS males had significantly lower odds of migraine (OR = 0.25, 95% CI = 0.20 – 0.32 and OR = 0.03, 95% CI = 0.02 – 0.04). However, contrary to our hypothesis, in comparison to SS females, OS females had significantly *higher* odds of migraine (OR = 1.51, 95% CI = 1.26 – 1.81). When SS males were included as the reference level, as expected, both SS and OS females demonstrated increased odds of migraine (OR = 4.08, 95% CI = 3.20 – 5.20 and OR = 6.15, 95% CI = 4.79 – 7.89), but OS males surprisingly had significantly *lower* odds of migraine (OR = 0.11, 95% CI = 0.08 – 0.15). See Figure 2 for odds ratios plot of migraine risk by twin type.

**Figure 2.**
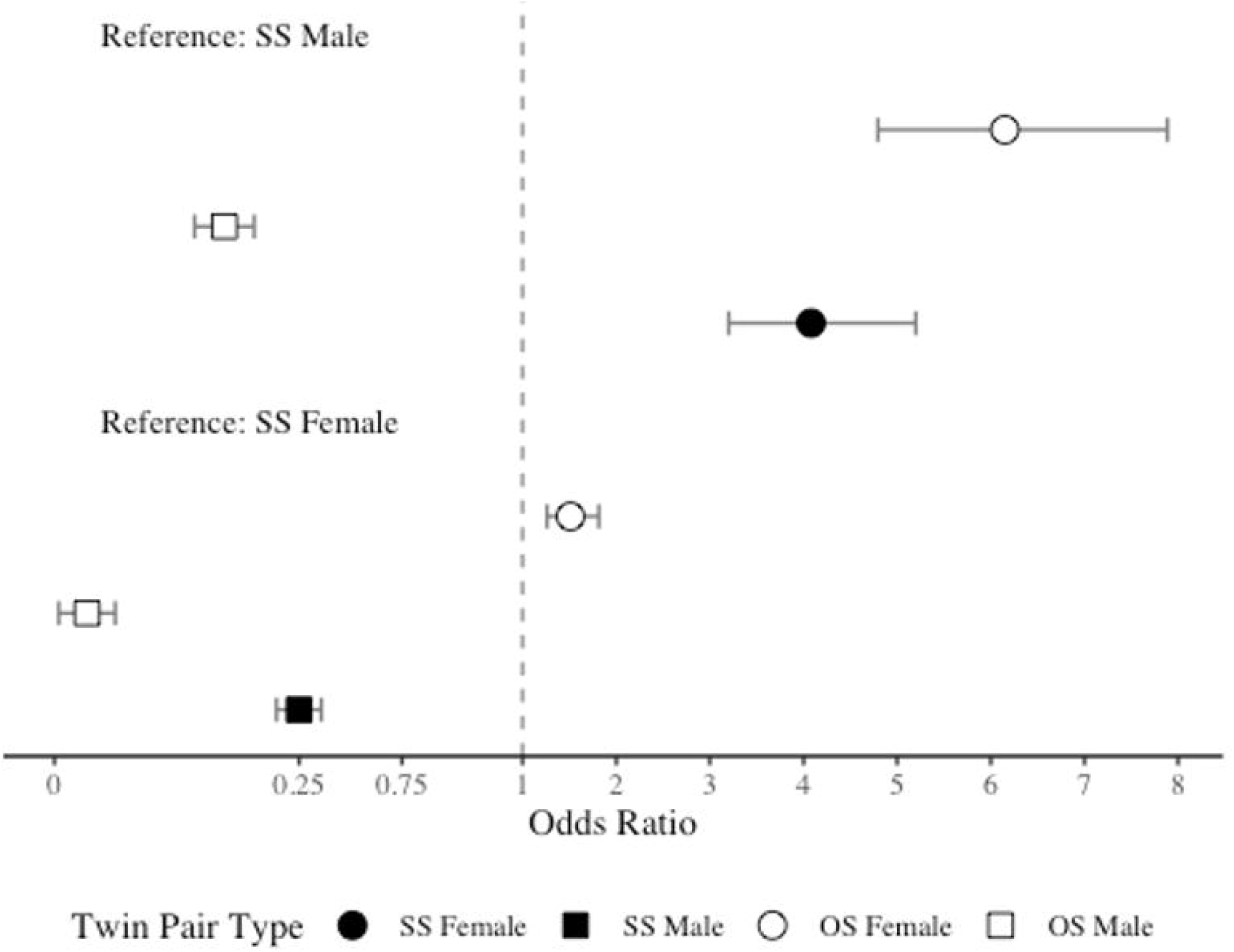
Odds ratio plot for migraine risk by twin type.

## 3 DISCUSSION

In the present study we utilized one of the largest single-cohort twin samples ever collected to examine the origins of sex differences in migraine. Our findings from the ADE sex-limitation model are largely in alignment with previous literature (Mulder et al., 2003; Gervil, Kaprio & Russell, 1999; Honkasalo et al., 1995). Specifically, we observed a broad-sense heritability of .45, which was primarily driven by non-additive genetic effects. Results from the ADE model also revealed that when additive and non-additive genetic influences were modeled together there is no evidence for qualitative or quantitative genetic differences in migraine risk across males and females. Prior studies, including those that have used data from the STR, have also not reported quantitative or qualitative genetic differences (Muller et al., 2003; Svensson et al., 2003). However, under a reduced model, one in which only additive genetic and unique environmental influences were modeled (AE model), we did find evidence for subtle *qualitative* differences in migraine across the sexes, while heritability remained equivalent.

Our finding of a qualitative genetic difference between men and women when a reduced, or simplified, genetic model was tested suggests that additional exploration of genetic differences in migraine is warranted. By fitting a model with only additive genetic influence, we reduced the number of model parameters and improved our ability to estimate the genetic correlation between OS twins. Thus, it may be the case that this effect is an artifact of the increased power of the analysis due to the simplification of the model. It is worth noting that the SALT and STAGE studies provided some of the most in-depth assessments of migraine available for genetically informative research. This in conjunction with the large sample size strongly supports the validity of the finding that non-additive and additive genetic factors contribute to migraine risk. Given this more complex genetic architecture of migraine, larger sample sizes will be needed to fully elucidate genetic effects, especially qualitative sex differences, both in the context of the twin design and genome-wide association studies.

Contrary to our hypothesis, the analysis of OS twins relative to SS female twins found that females with a male co-twin demonstrated a higher prevalence of migraine compared to females with a female co-twin. This result adds to a growing body of twin studies of behavioral phenotypes, perception, personality, cognition, and disease risk that support the ‘twin testosterone transfer hypothesis’ (Tapp, Maybery & Whitehouse, 2011; Loehlin & Martin, 2000; Verweij, Mosing, Ullén, Madison, 2016). However, this effect was in opposition to our predictions, as we theorized that the presence of a male in utero would decrease a female’s risk of migraine. The direction of the reported effect was surprising given that males have lower rates of migraine, suggesting that testosterone promotes resistance to the disease or demonstrates anti-nociceptive effects (Fischer et al., 2007; Vetvik & MacGregor, 2017; Shields, Seifert, Shelton, & Plato, 2019; van Oosterhout et al., 2018; Glaser, Dimitrakakis, Trimble, & Martin, 2012).

Although these findings were in opposition to our hypothesis, there is physiological evidence to suggest that the result are valid. For example, it is possible that female fetuses could have the biological mechanisms necessary to mitigate any lasting effects of exogenous gonadal hormones. In the developing female fetus, maternal testosterone is converted to estrogens via aromatase (Kallak et al., 2017). Thus, one theory is that the testosterone transferred to the female fetus from a male co-twin may be fully converted to estrogen before any lasting androgenization occurs. While this could explain why we did not observe a decrease in disease risk for females with a male co-twin, this would not account for the *increased* migraine risk observed for those females.

To address this point, we theorize that having a male co-twin could modify the female endocrine system such that females become more sensitive to estrogen, or changes in estrogen, throughout their lifetime. Evidence supports organizational effects of prenatal androgens on the female endocrine system including estrogen receptors (Landers, 2020; Kühnemann et al., 1994; Vito & Fox, 1981). It is also possible that prenatal hormones alter the female nervous system at the epigenetic level. Specifically, evidence suggests that in utero androgens impart lasting DNA methylation modifications in females with a male co-twin (Kong et al., 2020). OS females are discernable from SS females with OS females demonstrating greater similarities to male DNA methylation patterns (Kong et al., 2020). Critically, DNA methylation differences between OS females and SS females were related to nervous system regulation, a physiological component that strongly relates to migraine (Shechter, Stewart, Silberstein, & Lipton, 2002). Further research is clearly required to investigate these organizational and epigenetic theories, and to identify a more detailed mechanism to account for the observed results.

### Limitations

The present study is not without its limitations. First, the STR does not offer data on migraine days per-month, a descriptor that is used to diagnoses subtypes of migraine (i.e., episodic, chronic, refractory, etc.). This prevented us from being able to parse out migraine without aura subtypes. Furthermore, the STR is also lacking data for age at onset of migraine, which would be advantageous when addressing the influence of lifetime hormonal events on sex differences in migraine. Data also lacked information on lifetime course of migraine. It is unknown if most of the suspected cases of migraine reflect lifelong conditions, more short-term events or newly onset symptoms. While the diagnostic assessments utilized were thorough, we were nevertheless unable to sufficiently address migraine with aura according to the ICHD-3 criteria, which more than 30% of all migraine patients experience. There is an ongoing debate about if and to what degree migraine with and without aura are distinct entities (Kincses et al., 2019; Vgontzas & Burch, 2018; Granziera et al., 2006; Russell, Rasmussen, Fenger & Olesen, 1996).

### Future Directions

Results from the present study indicate that the sex-skew in the prevalence of migraine is not immediately attributable to differences in underlying genetic determinants. However, the presence of non-additive genetic effects complicates any genetically informed analysis, as most are based on simplified additive-only models. Even larger sample sizes than what was used here are needed to achieve sufficient power to fully elucidate the impact of these non-additive effects. Although the mechanism of action remains unclear, differences in the hormonal milieu of the prenatal environment do appear to alter disease risk for men and women. Under the traditional biometrical model of disease risk, the next avenue of exploration is likely to be interactions between genetic and environmental factors, as well as possible interactions between genetic and endocrine factors.

Ample knowledge is still to be gained surrounding the genetic epidemiology of migraine from data involving age at onset of disease and timing of critical hormonal events. These data will enable a thorough investigation into the genetic and environmental contributors to sex differences in migraine. The traditional approach of covarying for sex in a sample population with migraine is ineffective and dismissible as it does not account for the possibility of different factors contributing to migraine in men and women. Another potentially interesting future direction could be the examination of migraine risk among daughters of women with polycystic ovarian syndrome, due to fetal exposure to elevated testosterone levels (Dumesic et al., 2020). Understanding migraine in a sex-specific manner is an important goal as it holds promise for improving clinical care, diagnostic abilities, and therapeutic interventions for both men and women.

### Conclusions

Here we present the most in-depth analysis of the factors that contribute to migraine risk known to date. Specifically, we leveraged the largest migraine sample populations (n = 51,872) and present the first evidence that genetic factors in migraine risk may be different across the sexes. Additionally, although we hypothesized that the presence of a male co-twin would impart a masculinizing effect and reduce the risk of migraine in females, our analysis revealed an increase in migraine risk for females with a male co-twin relative to females with a female co-twin. Although the present analysis was unable to identify specific genes that differed across the sexes, we propose a series of potential mechanisms through which early environmental factors could influence migraine risk.

---

| Glossary of Terms |  |
| --- | --- |
| -2LL | Negative 2 log-likelihood |
| AIC | Akaike's information criterion |
| CI | Confidence Interval |
| DZ | Dizygotic |
| ICHD-3 | IHS International Classification of Headache Disorders version 3 |
| IHS | International Headache Society |
| LRT | Likelihood-ratio chi-square test |
| MO | Migraine Without Aura |
| MZ | Monozygotic |
| OR | Odds Ratio |
| OS | Opposite-Sex |
| SALT | Screening Across the Lifespan Twin Study |
| SS | Same-Sex |
| STAGE | Study of Twin Adults – Genes and Environment |
| STR | Swedish Twin Registry |

## 4 Conflict of Interest

The authors declare that the research was conducted in the absence of any commercial or financial relationships that could be construed as a potential conflict of interest.

## 5 Author Contributions

Morgan Fitzgerald and Matthew Panizzon are credited with the conception of the study. All authors contributed to data analysis and interpretation. Morgan Fitzgerald was the primary manuscript author. All authors contributed to revising and editing.

## 6 Funding

No grant funds were utilized.

## Notes

### Competing Interest Statement

The authors have declared no competing interest.

